# ZNF865 Regulates Senescence and Cell Cycle for Applications to Cell Engineering and Gene Therapy

**DOI:** 10.1101/2023.10.25.563801

**Authors:** Hunter Levis, Christian Lewis, Elise Stockham, Jacob Weston, Ameerah Lawal, Brandon Lawrence, Sarah E. Gullbrand, Robby D. Bowles

## Abstract

Zinc finger (ZNF) proteins represent the largest group of regulatory proteins within eukaryotic genomes. However, despite their broad regulatory function, the majority of ZNF protein function remains unknown. Recently, we discovered ZNF865, which has no in-depth publications and has not been functionally characterized. Utilizing CRISPR-guided gene modulation, we show that ZNF865 regulates key cellular and molecular processes associated with healthy cell function by primarily regulating cellular senescence, cell cycle progression, and protein processing. As a result, regulating this gene acts as a primary titratable regulator of cell activity, and we demonstrate the potential of targeted ZNF865 regulation as a tool to control senescence and protein production in multiple clinically relevant cell types for cell engineering/tissue engineering/gene therapy applications. We demonstrate its ability to rescue human senescent cell populations, boost T-cell activity, and dramatically deposit more cartilaginous tissue in a whole organ tissue-engineered intervertebral disc. Overall, we present novel biology and regulatory mechanisms of senescence and cell cycle that were previously unknown and display the power of CRISPR-cell engineering to enhance cell engineering strategies treating disease.

## Introduction

Zinc-finger (ZNF) proteins represent the largest group of regulatory proteins within eukaryotic genomes. Existing in nearly 5% of proteins, ZNF’s coordinate one or more zinc ions within the protein to stabilize ZNF structure, protein folding, and mediating interactions with DNA, RNA, lipids, and post-translational modifications^1–3^. Due to their extensive interactions, ZNFs regulate diverse cellular processes, such as transcription, cell migration, growth, proliferation, and differentiation^3,4^. Despite their assorted and distinct regulation of key cellular and molecular processes, the majority of ZNF protein function remains unknown^5^. We recently discovered a novel ZNF, ZNF865, that has never been functionally characterized before, yet regulates key cellular and molecular processes.

ZNF865 is a Cys_2_His_2_ (C2H2) ZNF that is 1059 amino acids long, containing 6 disordered regions, 20 unique ZNF domains, two transactivation domains (TADs), and two TGEKP linkers (**Figure 1A**)^6^. The presence of TADs indicates that ZNF865 likely stimulates transcriptional activity through contact with transcription factors (TFs) and associated components^7^. Approximately half of all C2H2 ZNFs contain a conserved TGEKP sequence which adopts a well-defined structure upon binding to DNA, linking adjacent fingers, and playing a vital role in DNA binding affinity^1,2^. Further, ZNF865 is broadly expressed across cell and tissue types, yet, there are no in-depth publications on ZNF865 indicating function (**Figure 1B**). Currently, a single manuscript referencing ZNF865 as a correlative biomarker in a panel of ZNF genes exists but contains no discussion of function or biological role of ZNF865^8^. To this end, our systematic investigation of ZNF865 will provide fundamental knowledge of biology and the cellular and molecular pathways regulated by ZNF865.

**Figure 1:**
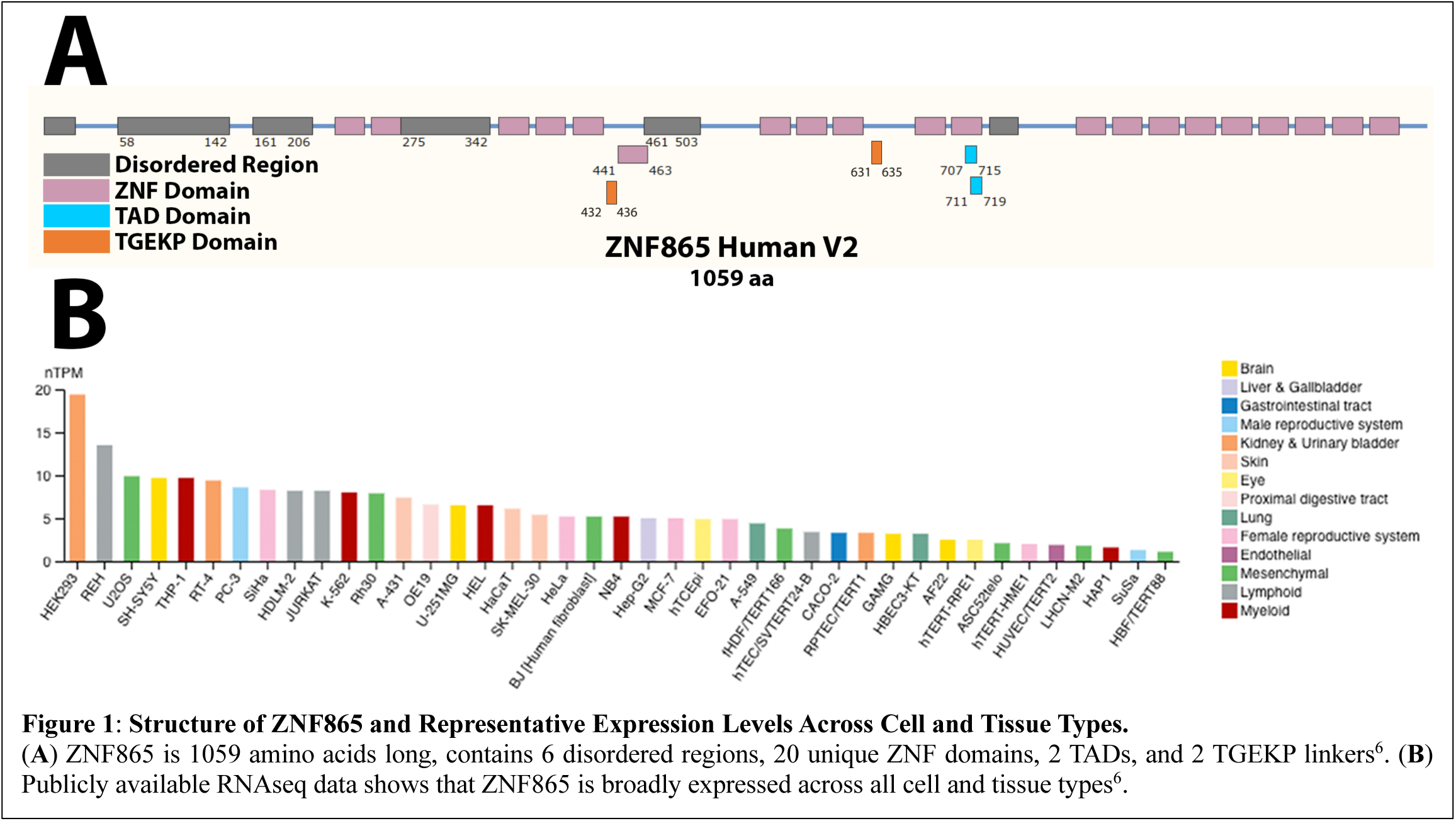
Structure of ZNF865 and Representative Expression Levels Across Cell and Tissue Types. (**A**) ZNF865 is 1059 amino acids long, contains 6 disordered regions, 20 unique ZNF domains, 2 TADs, and 2 TGEKP linkers6. (**B**) Publicly available RNAseq data shows that ZNF865 is broadly expressed across all cell and tissue types6.

Recently, clustered regularly interspaced short palindromic repeats (CRISPR)-guided gene modulation systems have emerged as powerful tools for profiling complex gene regulatory networks^9^, gene function^10^, improving cell survival^11^, and directing stem cell fate^12^. These systems comprise a single-guide RNA (gRNA) that targets the system to a gene of interest, and a nuclease-null mutant Cas9 (dCas9) that is fused to an effector molecule (VPR^13^ or KRAB^14^) for targeted upregulation (CRISPR-activation) or downregulation (CRISPR-interference). Using these systems allows for highly specific and titratable regulation of gene expression, both up and down. By modulating the expression of ZNF865, we can investigate the basic function and regulatory mechanisms of ZNF865 within the human genome and probe the highly complex and unknown facets of ZNF protein function.

Herein we present the initial characterization of ZNF865 utilizing CRISPR-guided gene modulation. Using RNAseq as our starting point, we correlated global changes in gene expression due to upregulation of ZNF865 to regulation of cellular senescence, cell cycle, DNA replication, mRNA transport, and protein processing across multiple unrelated cell types. To verify these correlations, we upregulated ZNF865 and showed increased rate of entry into the cell cycle, increased proliferation rates, and mRNA/protein processing rates. The suppression of ZNF865 leads to rapid induction of cellular senescence in primary cells and cell death in immortalized cells, indicating that expression of ZNF865 is necessary for normal cell function. Furthermore, we explore the ability of targeted ZNF865 upregulation to rescue cell populations from senescence and show that ZNF865 upregulation increases protein production and deposition rates in clinically relevant cell types with the goal of enhancing the clinical outcomes of cell/tissue engineering and gene therapies. Finally, a tissue engineered intervertebral disc (DAPS^15,16^) was used as a whole-organ model system to evaluate the ability of our ZNF865 CRISPR-engineered cells to produce functional physiological tissue in a promising tissue engineering strategy to treat degenerative disc disease (DDD). Overall, we demonstrate novel biology on ZNF865s critical role in cell function and potential application in cell engineering for cell therapies, gene therapies, and tissue engineering across a wide range of human diseases.

## Results/Discussion

### ZNF865 Upregulation Alters Similar Related Cell Functions Across Multiple Cell Types

RNAseq evaluated the global changes in gene expression due to upregulation of ZNF865 in three distinctly different cell types: HEK 293 cells, human adipose-derived stem cells (ASCs), and ACAN/COL2A1 CRISPRa upregulated ASCs (**Figure 2A**). Upregulation of ZNF865 in HEK 293 cells, ASCs, and ACAN/COL2A1 ASCs significantly differentially expresses 2,699, 6,647, and 8,093 genes, respectively. Overall, increased expression of ZNF865 differentially expressed thousands of genes across several cell types indicating broad regulatory control of molecular processes and gene expression.

**Figure 2:**
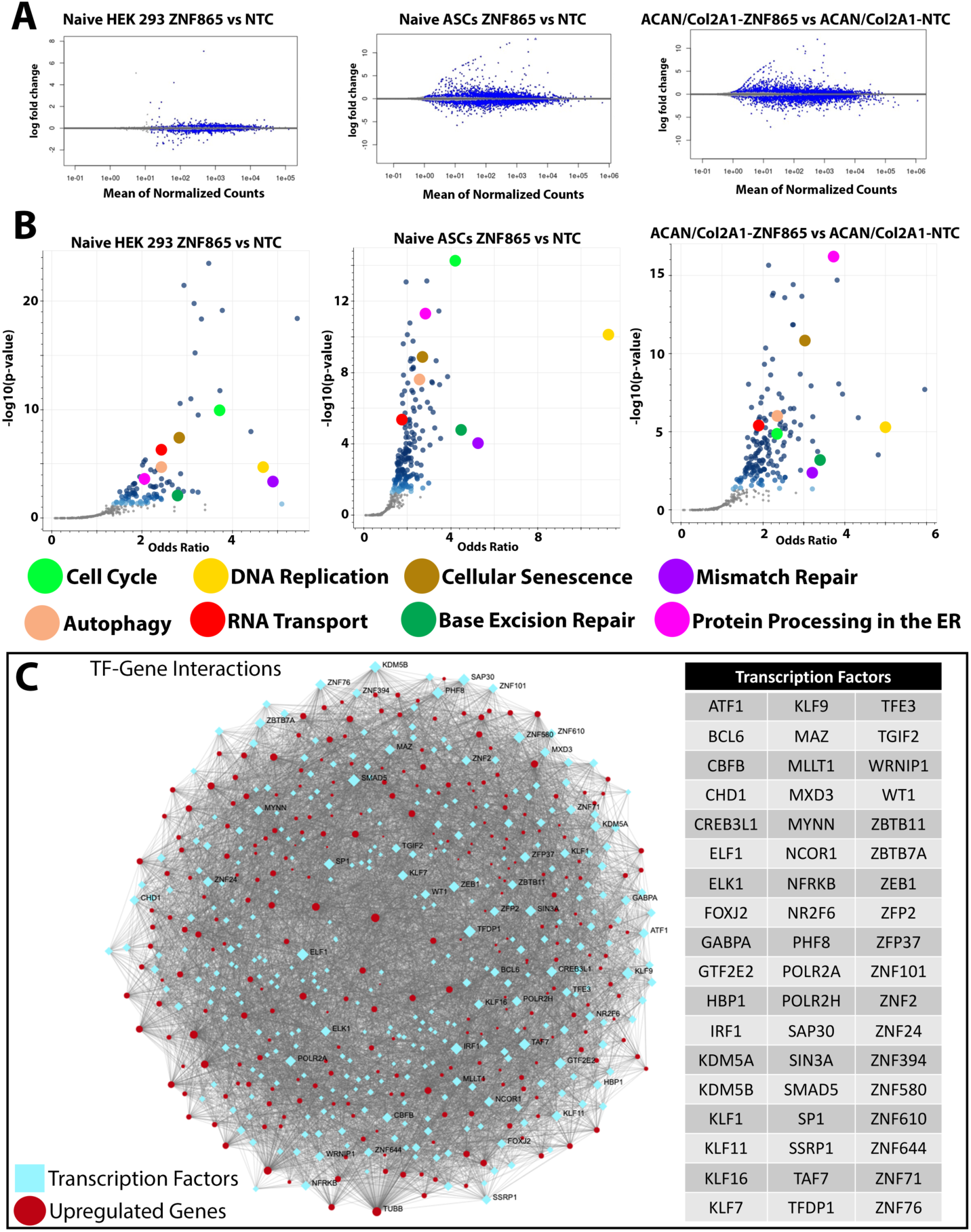
ZNF865 Regulates Key Cellular and Molecular Processes. RNA-seq on ZNF865 upregulated cells shows thousands of differentially expressed genes. (**A**) RNAseq plots for Naïve HEK 293 Cells, Naïve ASCs, and ACAN/COL2A1 ASCs show the number of genes significantly differentially expressed by increased ZNF865 expression. (**B**) Gene ontology analysis shows the top molecular processes affected due to increased ZNF865 expression in Naïve HEK 293, Naïve ASCs, and ACAN/COL2A1 ASCs. The 8 common top molecular processes affected by ZNF865 upregulation across cell types includes cell cycle, DNA replication, cellular senescence, mismatch repair, autophagy, RNA transport, base excision repair, and protein processing in the ER. (**C**) Transcription Factor-Gene network analysis shows the complex and highly interconnected network occurring due to upregulation of ZNF865. *NTC is nontarget control*.

Further analysis of the RNAseq data displays representative plots of the significantly differentially regulated molecular processes affected by ZNF865 upregulation. Utilizing the KEGG 2021 Human Pathway^17^, the top molecular processes affected by ZNF865 upregulation for all tested cell types are: cellular senescence, cell cycle, DNA replication, mismatch repair, autophagy, RNA transport, base excision repair, and protein processing in the ER (**Figure 2B**). Additionally, predictive binding analysis^18^ of the different ZNF binding motifs identified three molecular processes most likely affected by this protein which included cellular senescence, DNA replication, and cell cycle (*Supplementary Table 1*), consistent with our RNAseq analysis. Additional analysis of transcription factor (TF)-gene network interactions was analyzed using NetworkAnalyst to evaluate the complex and highly interconnected relationships of the significantly differentially expressed genes (DEGs) due to ZNF865 upregulation^19,20^. Overall, thousands of interactions are occurring due to increased expression of ZNF865 with one of the most significant TF’s being RNA polymerase II (POLR2A) (**Figure 2C**), which is predicted to interact with ZNF865^6,21^. Additionally, this list of TFs provides evidence for ZNF865 being a powerful regulator of gene expression within the human genome. Based on these results, we studied and verified ZNF865’s regulation of a subset of the top 8 molecular processes: cell cycle, DNA replication, and cellular senescence.

### ZNF865 Upregulation Regulates Cell Cycle Progression

Utilizing the CRISPRa system we can precisely target and upregulate ZNF865 (**Figure 3A/B**) in HEK 293 and ASCs to analyze shifts in the cell cycle through flow cytometry (**Figure 3C**). qRT-PCR verified ZNF865 upregulation, with 5 of 6 gRNA’s showing significant increases in ZNF865 expression. We can design guides targeting different regions of the genes promoter to titrate ZNF865 expression, with our top gRNA reaching a 7.8-fold increase in ZNF865 expression compared to the -NTC baseline expression levels of ZNF865 expression in ASCs (**Figure 3D**). A correlative analysis between average proliferation rates and fold change expression displays a strong correlation between ZNF865 expression and rates of cell proliferation in ASCs, displaying our ability to design gRNAs for precise control over cell activity and expression of ZNF865 (**Figure 3E**). Representative histograms display the shift in cell cycle and significant increases observed in proliferation rates for both HEK 293 (**Figure 3F**) and ASCs (**Figure 3G**). Analysis of the shift in cell cycle shows the percentage of cells in each phase of the cell cycle, with a significant shift of cells from G0/G1 into the S/G2/M phase in both ZNF865-edited HEK 293 cells (**Figure 3F**) and ASCs (**Figure 3G**). In addition to the shift in cell cycle, we observe significant increases in proliferation rates for HEK 293 cells and ASCs utilizing our top-performing gRNA (Guide 1). Both cell types show decreased doubling times due to ZNF865 upregulation where HEK 293 doubling time drops from 28.3 hours to 24.7 hours in ZNF865-edited HEK 293 cells and decreasing from 41.5 hours to 31.5 hours in ZNF865-edited ASCs (**Figure 3F**). The ability to decrease the doubling time and improve the expansion rate of cells is of broad clinical interest with cell therapies^22^. Overall, the targeted upregulation of ZNF865 could be used as a tool for controlling the expansion rate of cell-based therapies and regulate proliferation more broadly for therapeutics across many disease states.

**Figure 3:**
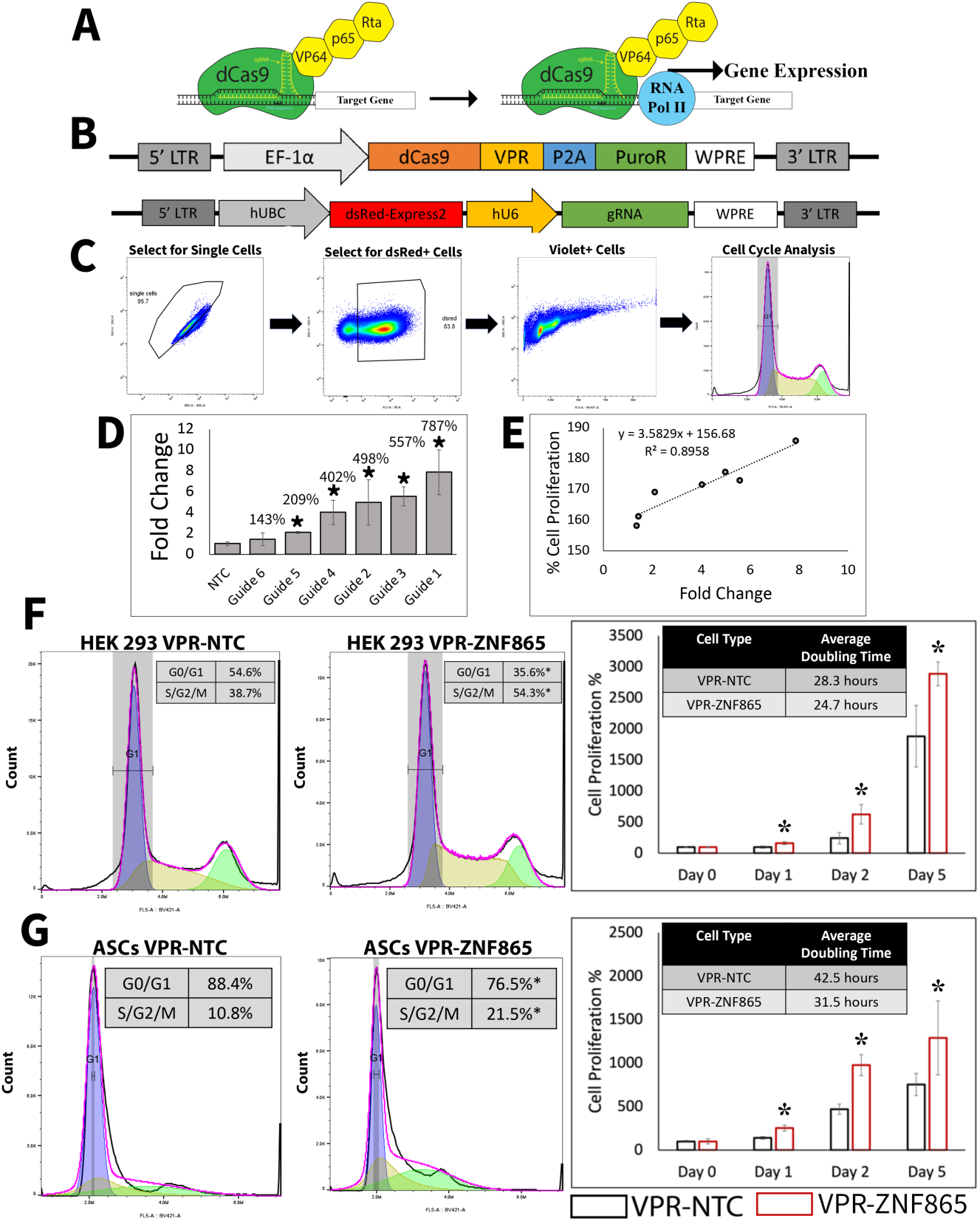
ZNF865 Regulates Cell Cycle Progression. Targeted upregulation of ZNF865 drives entry into the cell cycle in HEK 293 cells and ASCs. (**A**) The CRISPRa system utilizing the VP64-p65-Rta (VPR) effector molecule acts like a synthetic transcription factor, recruiting RNA polymerase II and effectively upregulating target gene expression. (**B**) The dCas9-VPR expression cassette and targeted gRNA expression cassette for ZNF865 or NTC upregulation. (**C**) Representative plots showing the gating strategy for cell cycle analysis. (**D**) qRTPCR displays our ability to upregulate and control expression of ZNF865 gRNA design in ASCs (n=4, *=p<0.05). (**E**) A correlation plot between the Fold Change in expression of ZNF865 and % Cell Proliferation displays a strong correlation between expression of ZNF865 and cell proliferation rate in ASCs (R2 = 0.8958). Increased ZNF865 expression shifts the percentage of cells in each stage of the cell cycle, with cells entering the cell cycle at a higher rate in the ZNF865-edited cells compared to the -NTC cells. Representative histograms show the shift in cell cycle between the -NTC and -ZNF865 edited as well as increased proliferation rates in **(F**) HEK 293 cells and (**G**) ASCs, and a decrease in average doubling rates of both cell types edited with ZNF865 compared to the NTC. (*=p<0.05, n=4). *NTC is nontarget control*.

### ZNF865 Expression is Necessary for Healthy Cell Function

The CRISPRi system can effectively suppress ZNF865 expression by tri-methylation of the histone at the targeted location of our designed gRNAs, preventing transcription of the target gene and suppressing gene expression (**Figure 4A**)^23^. Quantitative reverse transcriptase polymerase chain reaction (qRT-PCR) verified downregulation of ZNF865 expression in ASCs, with 4 of 5 guides displaying statistically significant decreases in expression and Guide 3 showing 93.3% suppression of ZNF865 (**Figure 4B**). The ability to highly control expression of ZNF865 displays the power of the CRISPRi system to tightly control gene expression for broad applications of cellular engineering.

**Figure 4:**
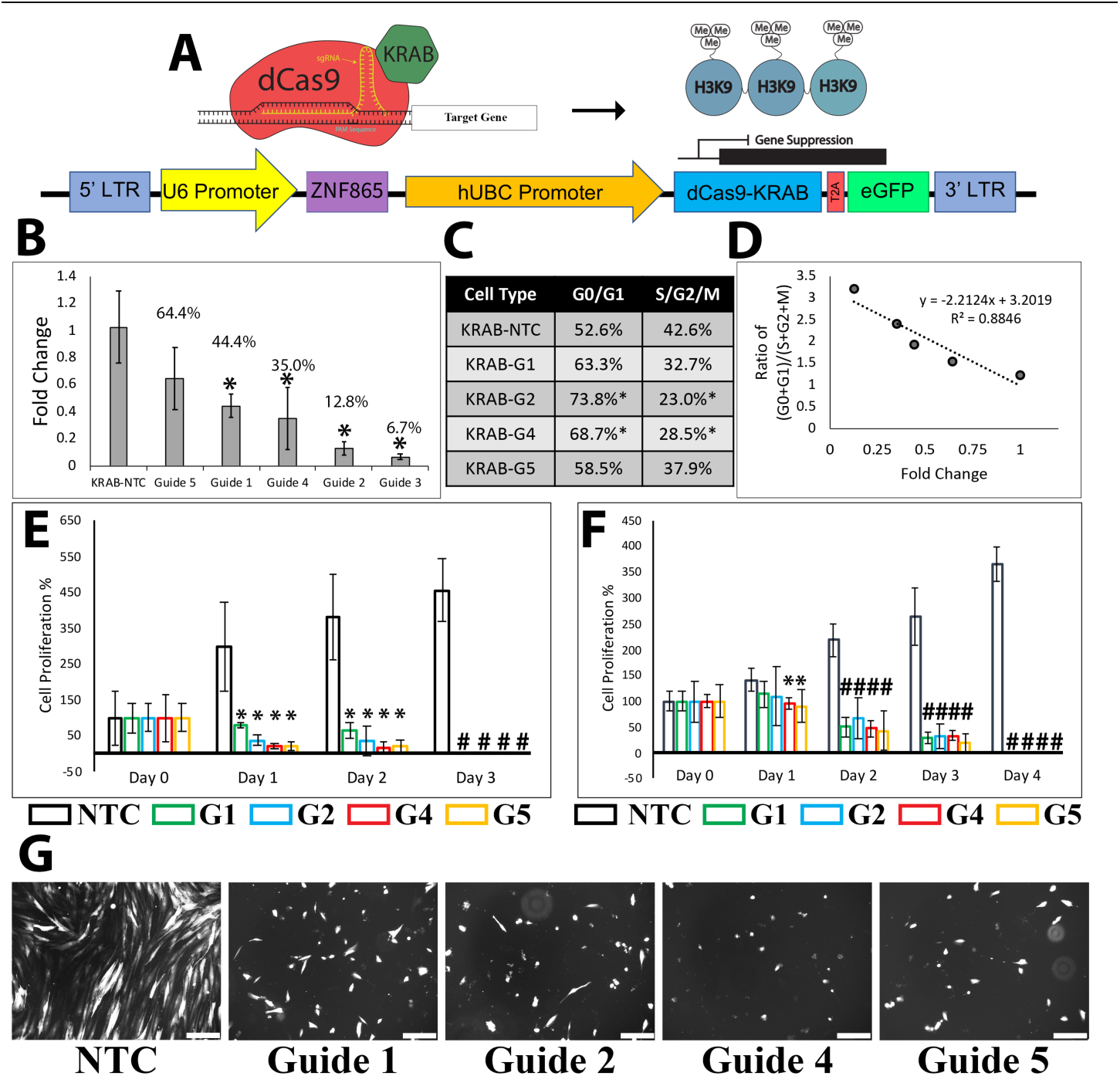
ZNF865 Expression is Necessary for Cell Survival. Targeted downregulation of ZNF865 using CRISPRi induces cell death in HEK 293 cells and ASCs. (**A**) The CRISPRi system utilizes the effector molecule KRAB which trimethylates the histone, preventing transcription and suppressing targeted gene expression. (**B**) qRT-PCR verifies downregulation of ZNF865, with guides 1, 4, 2, and 3 significantly suppressing ZNF865 expression, percent expression is displayed on the plot. (**C**) Downregulation of ZNF865 leads to prevention of HEK 293 cells entering the cell cycle with a buildup of cells in the G0/G1 phase. (**D**) Correlation analysis of fold-change and ratio of percent HEK 293 cells in (G0+G1)/(S+G2+M) phase of the cell cycle showing a strong correlation between expression of ZNF865 and cell cycle progression (R2=0.8846). Downregulation of ZNF865 in (**E**) HEK 293 cells and (**F**) ASCs induces to cell death within 3 or 4 days, respectively (*=p<0.05, #=p<0.001, n=4). (**G**) Representative images showing ASC morphology on Day 2 (scale bar = 200μm). *NTC is nontarget control*.

Cell cycle analysis of HEK 293 cells 96-hours after transduction shows a noticeable shift in the cell cycle. Compared to upregulation where we observed cell cycle shifting towards mitosis, when ZNF865 is downregulated, we observe an increase in population of cells within the G0/G1 phase. Guides 2 and 4 show a significant shift in the cell cycle with nearly 74% of the cells in Guide 2 being within the G0/G1 phase (**Figure 4C**). This trend continues for the other guides with a noticeable, but nonsignificant, shift in the cell cycle. When we correlate the cell cycle data with fold-change in expression we observe a strong correlation between shifts in cell cycle and expression of ZNF865 (R^2^=0.8846, **Figure 4D**). Furthermore, when we analyze cell proliferation rates after successful transduction with the CRISPRi system we observe cell death in HEK 293 cells (**Figure 4E**) and ASCs (**Figure 4F**) within 3 or 4 days after expression of the CRISPRi system. Representative images display ASC morphology and overall cell numbers 96-hours post-transduction, with all guides showing varying degrees of cell death (**Figure G**). Noticeably, Guide 3, which suppressed expression by 93.3%, is not shown on these plots because cell death was observed within 72-hours post transduction. Subsequent experiments evaluated the differential expression of ZNF865 in human primary nucleus pulposus cells (hNPCs) using the CRISPRi and CRISPRa system, to evaluate ZNF865’s role in inducing cellular senescence or rescuing primary cell populations from cellular senescence. Overall, ZNF865 is broadly expressed across cell types, and we wanted to evaluate the role of ZNF865 expression in hNPCs.

### ZNF865 Regulates Cellular Senescence in Primary Cells

Low-back pain and DDD are commonly linked to cellular senescence within the nucleus pulposus (NP) of the intervertebral disc (IVD)^24^. To evaluate ZNF865’s role in inducing cellular senescence healthy primary hNPCs were isolated from discarded surgical tissue from trauma patients and transduced with the ZNF865 downregulation expression cassettes (Guide 2 and Guide 3). Following successful transduction, hNPCs were monitored for cell proliferation over the course of 6-days and showed no proliferation compared to NTC hNPCs (**Figure 5A)**. After 1 and 3-weeks of culture, hNPCs were stained for key markers of cellular senescence, senescence associated-β-galactosidase (β-gal), p16^INK4a^ (p16), and p21^CIP^^1^ (p21)^25–27^. Quantified β-gal staining shows significant increases in positively stained cells after 1-week and 3-weeks of culture, with 76.8% of cell staining positively for β-gal at 3-weeks for Guide 3 (**Figure 5B**). Furthermore, p16 and p21, two key regulators of cell cycle progression and markers indicating cellular senescence, were used to further confirm ZNF865’s regulation of cellular senescence^25–27^. Quantified immunofluorescence after 3-weeks of culture shows a similar trend, with significant increases in staining for p16 (**Figure 5C**) and p21 (**Figure 5D**) in ZNF865-edited cells compared to the NTC. Representative images show the qualitative β-gal, p16, and p21 staining at 1 and 3-weeks of culture with noticeable increases in staining for Guide 3 ZNF865-edited hNPCs indicating the induction of cellular senescence (**Figure 5E**). Downregulation of ZNF865 confirmed ZNF865’s role as a regulator of cellular senescence, as originally suggested by the RNAseq data. By monitoring cell proliferation rates and evaluating the presence of key cellular senescence markers in hNPCs, our data confirms ZNF865’s role in regulating senescence, indicating that ZNF865 expression is necessary for healthy cell function.

**Figure 5:**
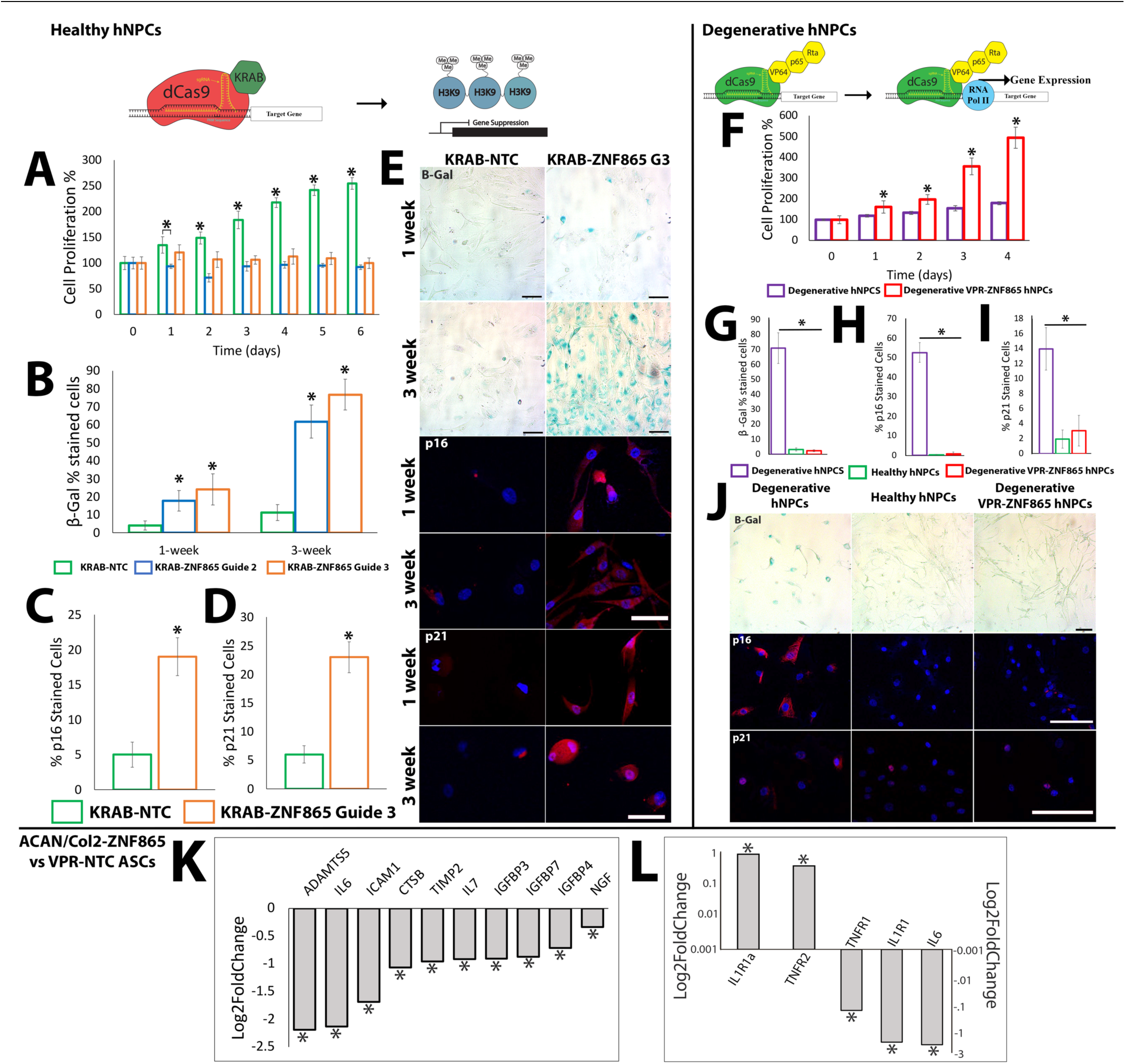
CRISPRi of ZNF865 Induces Senescence in hNPCs while CRISPRa of ZNF865 Rescues Populations of Degenerative hNPCs from Senescence. Targeted modulation of ZNF865 in hNPCs leads to the induction of senescence when ZNF865 expression is suppressed and rescues degenerative hNPCs from senescence when upregulated. (**A**) Cell proliferation of hNPCs after transduction with KRAB-ZNF865 Guide 2 or Guide 3 shows no cell proliferation compared to the KRAB-NTC (n=4, *=p<0.05). (**B**) Quantified β-gal staining shows increased staining for both guides after 1 and 3-weeks of culture, with Guide 3 showing nearly 80% of cells showing β-gal staining at 3-weeks (*=p<0.05, n=4). Quantified percent-stained cells for (**C**) p16 and (**D**) p21 3-weeks after transduction (n=4, *=p<0.05). Representative images of (**E**) β-gal staining (scale bar = 100μm), p16, and p21 immunofluorescence showing increased staining at 1 and 3-weeks of culture for Guide 3 compared to the NTC indicating ZNF865 is a regulator of cellular senescence. (**F**) Targeted upregulation of ZNF865 in degenerative hNPCs shows increased proliferation rates compared to unedited degenerative hNPCS (*=p<0.05, n=4). Quantified staining for (**G**) β-gal, (**H**) p16, and (**I**) p21, showed significant increases in staining for the degenerative hNPC control group and minimal staining in both the healthy hNPC group and the degenerative ZNF865-edited hNPC group (n=4, *=p<0.05). (**J**) Representative images show the β-gal staining, p16, and p21 immunofluorescence for degenerative hNPCs, healthy hNPCs, and degenerative VPR-ZNF865-edited hNPCs (scale bars = 100μm). Additional analysis of RNAseq data shows that ZNF865 significantly differentially regulates (**K**) genes associated with SASP and (**L**) genes associated with anabolism (IL1R1a and TNFR2) and catabolism (TNFR1, IL1R1, and IL6) of the IVD. *NTC is nontarget*.

### ZNF865 Upregulation in Degenerative hNPCs Rescues Cell Populations from Senescence

Degenerative primary hNPCs were isolated from discarded surgical tissue, from patients undergoing discectomy, and transduced with the CRISPRa system for targeted upregulation of ZNF865. Following successful transduction, degenerative hNPCs expressing the VPR-ZNF865 CRISPRa system were monitored for proliferation compared to the degenerative control. Targeted upregulation of ZNF865 in degenerative hNPCs significantly increases proliferation rates compared to degenerative hNPCs (**Figure 5F**). Additionally, ZNF865 decreases the average doubling time for degenerative hNPCs, dropping their average doubling time from 108 hours to 42 hours. Subsequently, degenerative hNPCs, healthy hNPCs, and degenerative VPR-ZNF865-edited hNPCs were fixed and evaluated for β-gal, p16, and p21 expression to confirm ZNF865-edited hNPCs were not expressing key markers related to senescence. Degenerative hNPCs showed significantly higher percentages of β-gal (**Figure 5G**), p16 (**Figure 5H**), and p21 (**Figure 5I**) expression compared to both the healthy hNPCs and the ZNF865-edited degenerative hNPCs. Furthermore, there was no significant difference in staining between the healthy hNPC group and the degenerative VPR-ZNF865 hNPC group indicating that ZNF865-edited degenerative hNPCs were not senescent. Representative images verify the expression of β-gal, p16, and p21 in degenerating hNPCs compared to healthy hNPCS and ZNF865-edited hNPCs, which show virtually no expression of these markers (**Figure 5J**). Furthermore, healthy hNPC morphology is conserved in the ZNF865-edited degenerative hNPC group, indicating that targeted upregulation of ZNF865 rescues hNPCs from senescence while not altering their morphological phenotype.

In addition, upregulation of ZNF865 results in significant downregulation of key genes implicated in the senescence-associated secretory phenotype (SASP)^28^ and in DDD^29^. ZNF865 significantly downregulates key genes associated with the SASP, including ADAMTS5, IL6, and NGF (**Figure 5K**). Further, key cytokines and receptors implicated in DDD (TNFR1, IL1R1, and IL6) are significantly downregulated while IL1Ra and TNFR2 are significantly upregulated and have been shown to provide a regenerative affect in DDD indicating that our ZNF865-edited ASCs may provide additional protective and regenerative effects *in vivo* (**Figure 5L**).

Senescent cells have been closely tied to a number of diseases and biological procecess such as aging^30^, osteoarthritis (OA)^31^, DDD^24^, cancer^30^, or Alzheimer’s^28,32^. As previosuly discussed and shown, we can control the induction of senescence (CRISPRi downregulation) and exiting of senesence (CRISPRa upregulation) in multiple unrelated cell types. The ability to highly control and regulate cells entering and exiting senescence would have dramatic implications for therapeutic application across a broad range of diseases. By rescuing cell populations from entering senescence by upregulating ZNF865 in degenerative hNPCs, we have shown promising results that could have profound implications on treating age-associated phenotypes and diseases that are often correlated with senescence^28^. However, the ability to control senescence by either inducing or rescuing cells from senescence may have severe consequences and it is paramount that a high degree of control and care is undertaken when regulating such a significant cellular process. Our results display a high degree of control over regulation of ZNF865 and it’s ability to induce or rescue cell populations from senescence. The continued investigation of this gene and its regulation of senescence will provide vital insight into the further development of cell and gene therapies that could treat many senescence-related diseases.

### ZNF865 Upregulation Amplifies Protein Production in Clinically Relevant Cell Types

To examine ZNF865’s affect in a broader range of cell types and to specifically test a cell type related to immune therapy, ZNF865 upregulation was performed in Jurkat cells and subsequently activated to evaluate cytokine secretion using an ELISA (**Figure 6A**). By increasing the proliferation rates and cytokine secretion rates of activated T-cells, we can show ZNF865’s ability to improve the expansion rate of T-cell therapies and the efficacy of immune therapies with increased cytokine secretion. In ZNF865-edited Jurkat cells we observed significantly increased proliferation rates compared to the NTC (**Figure 6B**). In addition, we observe significant increases in IL-2 (**Figure 6C**) and IFN-γ (**Figure 6D**) secretion in ZNF865-edited cells compared to NTC-edited cells after activation. There is a 2x increase in proliferation rate and 1.5x increase in production of IL-2 and IFN-γ in ZNF865-edited T-cells, indicating that targeted upregulation of ZNF865 can improve the expansion rate and efficacy of cancer cell therapies^33^.

**Figure 6:**
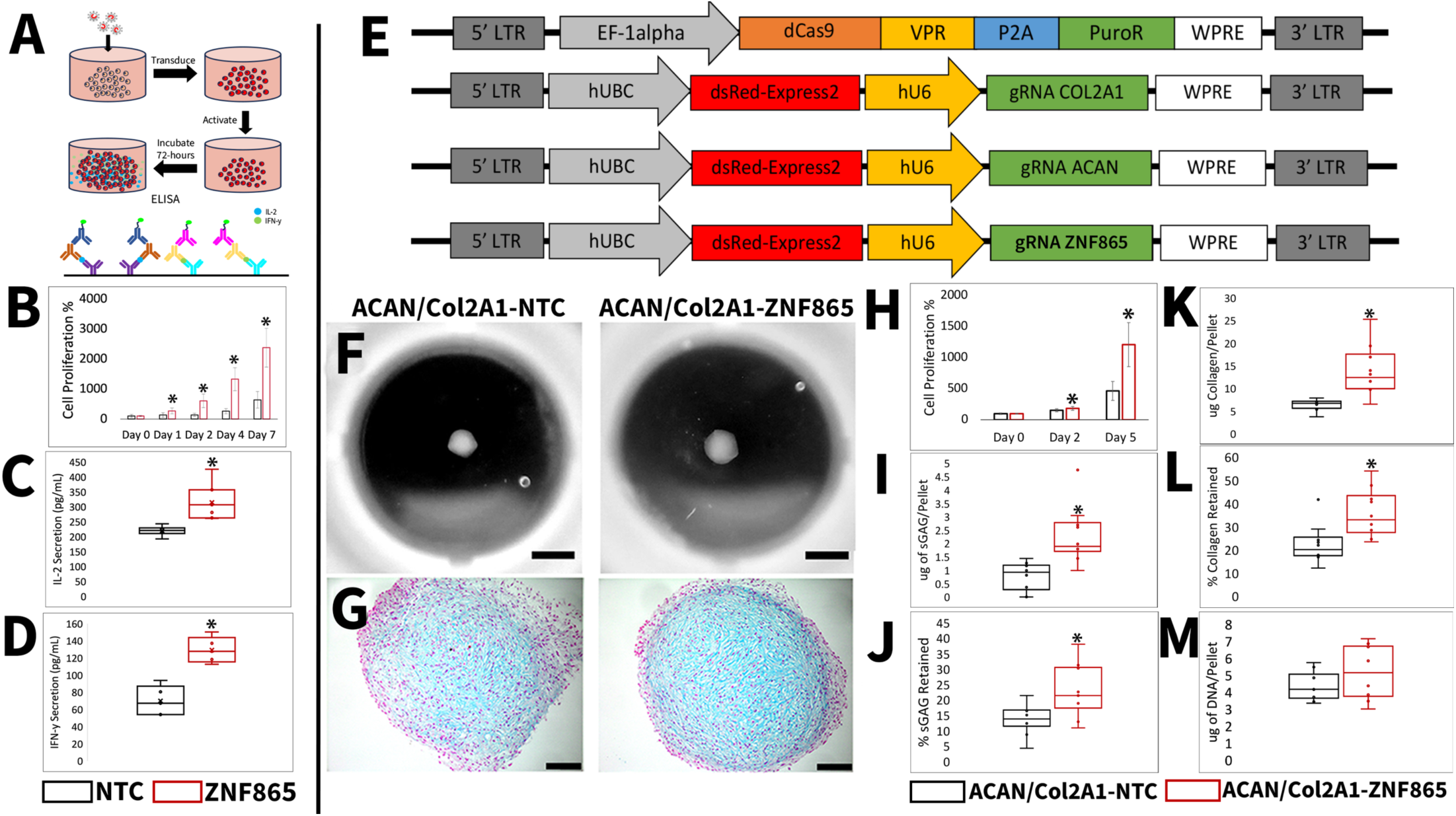
CRISPRa of ZNF865 Amplifies Cell Phenotype. Targeted Upregulation of ZNF865 Increases the Protein Processing Rates and Amplifies Cell Phenotype in Jurkat cells and ACAN/COL2A1 ASCs. (**A**) Workflow schematic of T-cell transduction, activation, and ELISA analysis. Jurkat-edited cells we observe a significant increase in (**B**) cell proliferation, and significant increases in (**C**) IL-2 and (**D**) IFN-y secretion after stimulation with Con A indicating enhanced protein processing and cytokine production due to -ZNF865 upregulation compared to the NTC. (**E**) The CRISPRa multiplex upregulation expression cassettes for the upregulation system and targeted genes COL2A1, ACAN, and ZNF865. ZNF865 upregulation qualitatively shows increased (**F**) tissue deposition (scale bar = 1mm) and (**G**) sGAG deposition (scale bar = 100μm). When expanding ACAN/COL2A1-edited cells in monolayer, (**H**) ZNF865-edited cells proliferate significantly faster compared to the ACAN/COL2A1-NTC. Biochemical analysis shows significant increases (**I**) concentration of sGAG per pellet and (**J**) retention of sGAG within the ZNF865-edited pellets. There are also significant increases in (**K**) collagen per pellet and (**L**) retention of collagen per pellet in ZNF865-edited pellets. There is no significant difference in (**M**) DNA content per pellet between the two groups (*=p<0.05, n=4-10). *NTC is nontarget control*.

Previously, our group demonstrated multiplex CRISPRa upregulation could drive a chondrogenic phenotype in ASCs by upregulating the genes aggrecan (ACAN) and collagen type II (COL2A1)^12^. These CRISPR-engineered ASCs have therapeutic applications in treating both osteoarthritis of the knee and DDD. We believed that multiplexing ZNF865 with ACAN and COL2A1 would lead to further increases in cartilage deposition compared to ACAN/COL2A1 alone. Therefore, 3D pellet culture was used to evaluate protein production and cartilaginous ECM deposition *in vitro* for ZNF865-edited ACAN/COL2A1 upregulated ASCs.

ZNF865 was multiplex upregulated in conjunction with COL2A1 and ACAN using the dCas9-VPR CRISPRa system (**Figure 6E)**. After 21-days of pellet culture ZNF865-edited pellets qualitatively showed increases in overall tissue deposition (**Figure 6F**) and sGAG deposition (**Figure 6G**) compared to ACAN/COL2A1-NTC cells. In monolayer, ACAN/COL2A1-ZNF865 ASCs proliferate significantly faster compared to ACAN/COL2A1-NTC cells (**Figure 6H**). Biochemical analysis shows ZNF865-edited cells produce significantly more sGAG/pellet (**Figure 6I**) and retain more sGAGs within the pellet compared to the NTC (**Figure 6J**). Collagen content shows the same trend, with significant increases in collagen/pellet (**Figure 6K**), and a significantly higher retention of collagen compared to the NTC (**Figure 6L**). Additionally, there is no significant difference in DNA content/pellet compared to NTC, yet there is twice as much sGAG and collagen produced in ZNF865-edited pellets compared to the NTC (**Figure 6M**). Taken together, our ability to improve immune cell cytokine secretion or cartilage ECM deposition displays broad functional applications in cell engineering and cell therapies and verifies the ability for ZNF865 to be combined with other cell engineering methods to boost their outcomes^40^.

### Multiplex ACAN/COL2A1-ZNF865 Upregulation Amplifies Cartilage Deposition in Engineered Disc

The intervertebral disc (IVD) is a fibrocartilaginous tissue comprised of three main parts: the cartilaginous endplates, nucleus pulposus (NP), and annulus fibrosus (AF), which allows for motion, weight bearing, and flexibility throughout the spinal column^34^. DDD is the breakdown of these IVDs, which is commonly linked to LBP, the leading cause of disability worldwide^35^. DDD is a complex multifactorial disease resulting in chronic inflammation and pain^34^. Recently, tissue engineered total artificial disc replacement (ADR), such as disc-like angle ply structure (DAPS)^15,16^, are being developed as better treatments compared to the current gold-standard, such as spinal fusion or discectomy^36^. DAPS is a tissue engineered ADR comprised of a cell-seeded agarose NP and an electrospun cell-seeded poly(ε-caprolactone) (PCL) AF where disc matrix synthesis is driven through growth factors and scaffold cues, which would allow for natural motion, physiologic weight bearing, and allow for normal flexion of the spine^15,16,37^. However, challenges with robust homogenous matrix distribution throughout the DAPS have been demonstrated with scale up to clinically relevant size scales during culture with chondrogenic growth factors^16^. Therefore, we hypothesized that our ACAN/COL2A1-ZNF865 CRISPR-engineered ASCs seeded in DAPS would produce similar or elevated levels of cartilage as is observed with stem cells dosed with growth factors. Further, our results would strongly indicate whether our CRISPR-engineered ASCs would perform favorably as a cell therapy treating DDD.

DAPS were seeded with ACAN/COL2A1-ZNF865 CRISPRa-edited ASCs to investigate matrix deposition in whole organ tissue engineering application. Naïve ASCs, multiplex CRISPRa ACAN/COL2A1, or multiplex ACAN/COL2A1-ZNF865 ASCs (**Figure 7A**) were seeded onto AF and NP regions of DAPS and cultured for a total of 5-weeks (**Figure 7B**). After 5-weeks of culture, the naïve ASCs were able to seed 7 total DAPS but only 1 DAPS was considered successful (i.e. the cells were able to produce sufficient matrix to adhere the AF scaffold layers into a single unit) after 5-weeks of culture, ACAN/COL2A1 ASCs were able to seed 6 DAPS with 3 successful DAPS, however our ACAN/COL2A1-ZNF865 cells were able to seed 9 total DAPS and all 9 were deemed successful (**Figure 7C**). Initially, our data confirms increased proliferation rates for ZNF865-edited DAPS compared to both control groups and the effectiveness of ZNF865-edited DAPS to produce ECM within DAPS. Alcian blue and picrosirius red combinatorial stain displays dramatic increases in tissue deposition for both the ACAN/COL2A1-ZNF865 ASCs compared to the ACAN/COL2A1-NTC ASCs (**Figure 7D**). Further analysis for cell phenotype using the RNAseq data displays the normalized counts for COL10A1, COL2A1, and ACAN ACAN/COL2A1-ZNF865 ASCs indicating that the chondrogenic phenotype is maintained after ZNF865 upregulation (**Figure 7E**). Individual stains further confirm our results with picrosirius red staining showing increases in collagen deposition in ACAN/COL2A1 and ACAN/COL2A1-ZNF865 DAPS compared to the naïve control (**Figure 7F**). Alcian blue staining shows the same trend, with increased staining for ACAN/COL2A1 and dramatically darker staining for ACAN/COL2A1-ZNF865 DAPS compared to the naïve control (**Figure 7G**). Moreover, our results display the effectiveness of targeted multiplex upregulation using CRISPRa for highly specific control of cell phenotype to amplify and drive a chondrogenic phenotype while simultaneously regulating the expression of the SASP in tissue engineered IVDs. Furthermore, we observe that ZNF865 can be used as a highly effective tool for enhancing cell therapies to treat DDD but also in DAPS to enhance tissue deposition and decrease the manufacturing rates of tissue engineered ADRs. Taken together, our ability to regulate protein production and senescence would be a powerful tool for gene, cell, and tissue engineering therapies.

**Figure 7:**
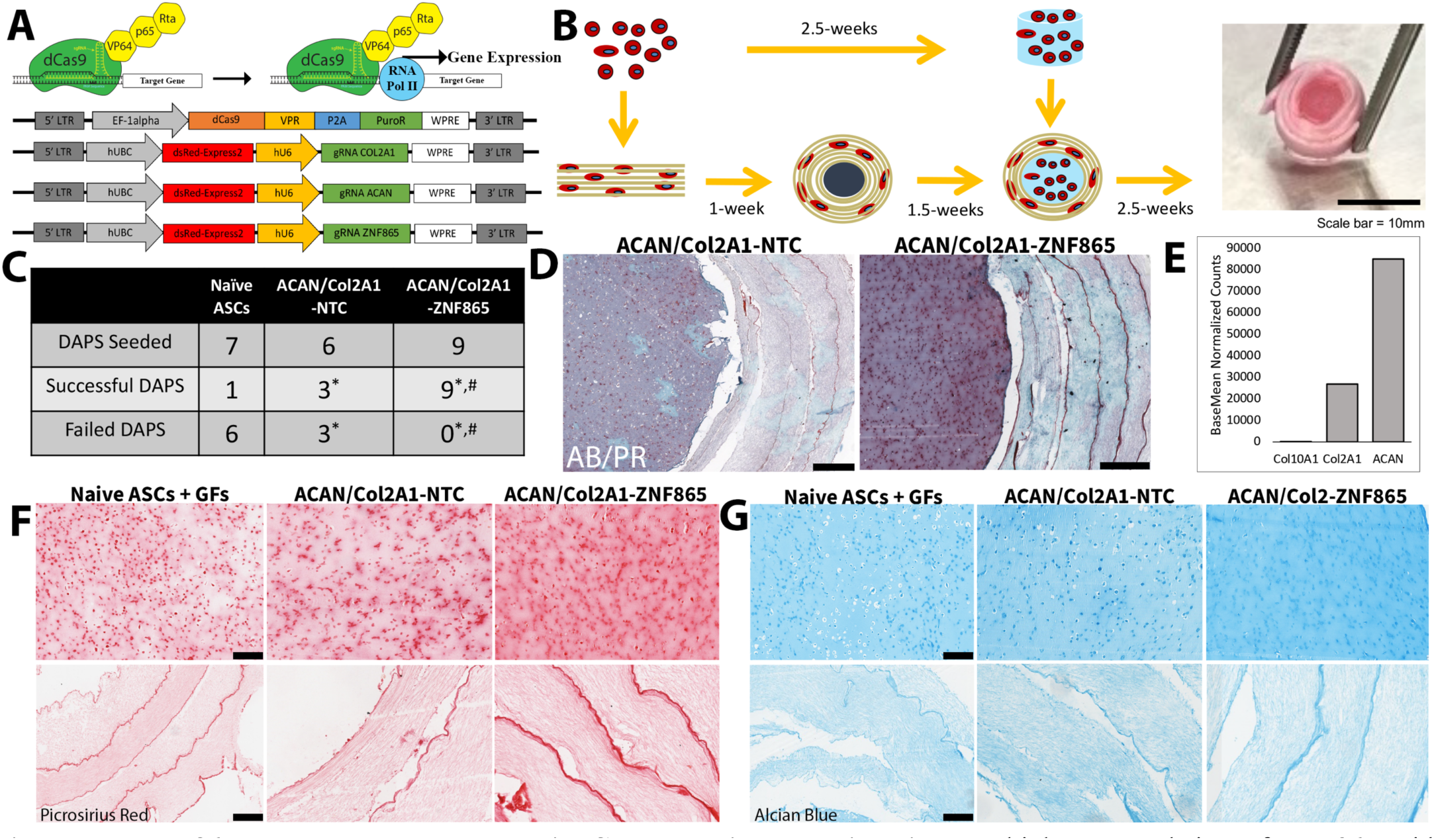
ZNF865 can be used as a Tool in Cell and Tissue Engineering. Multiplex upregulation of ZNF865 with ACAN/COL2A1 dramatically increases cartilage deposition in DAPS. (**A**) Representative schematic of the CRISPRa system and expression cassettes showing increased expression and targeted upregulation of ACAN, COL2A1, and ZNF865. (**B**) Workflow schematic of cell seeding and culture of DAPS over 5-weeks of total culture. (**C**) Representative Alcian blue/picrosirius red dual stains showing dramatic increases in ECM deposition between the naïve ASCs, ACAN/COL2A1 upregulation, and ACAN/COL2A1-ZNF865 upregulation (scale bar = 500um). (**D**). Over the course of 5-weeks only 1 successful DAPS was formed with the naïve ASCs, 3 with the ACAN/COL2A1 ASCs, and 9 with the ACAN/COL2A1-ZNF865 ASCs. (**E**) Average counts are shown for ACAN, COL2A1, and Col10A1 showing the chondrogenic phenotype is maintained in ZNF865-edited ACAN/COL2A1-ASCs. Representative (**F**) picrosirius red staining and (**G**) Alcian blue staining showing the NP and AF regions of the engineered disc showing dramatically more cartilage and sGAG deposition in the ACAN/COL2A1-ZNF865 ASCs compared to the Naïve and ACAN/COL2A1 ASCs (scale bar = 200μm).

## Outlook

We have reported the initial characterization of a novel gene, ZNF865 and displayed the application of ZNF865 as a tool to enhance cell and tissue engineering therapies. This research provides novel biology and regulatory mechanisms of key cellular and molecular processes that were previously unknown. Furthermore, this work demonstrates that targeted regulation of ZNF865 can control cellular senescence and amplify cell phenotype that can be used clinically to treat a wide range of human disease, including, but not limited to, OA of the knee or disc degeneration, two of the leading causes of disability worldwide^35^. In addition, the proof-of-concept study in DAPS^16^ displays the effectiveness of our ACAN/COL2A1/ZNF865-edited ASCs to produce cartilage while simultaneously downregulating genes associated with DDD and SASP. Overall, the ability to highly control the expression of ZNF865, rescue cell populations from senescence, and increase the proliferation and protein production rates of cells has broad applications across clinical applications and the continued research and understanding on ZNF865’s role in the human genome can provide valuable insight into developing novel gene and cell therapies.

## Supporting information

Supplemental RNAseq File

Supplemental Predictive Binding Data (Table 1 included)

## Acknowledgements

Sequencing was performed at the DNA Sequencing Core Facility, University of Utah. Flow cytometry and FACS was supported by the Office of The Director of the National Institutes of Health under Award Number S10OD026959 and NIC Award Number 5P30CA042014-24. Research reported in this publication utilized the High-Throughput Genomics and Cancer Bioinformatics Shared Resource and the Biorepository and Molecular Pathology Shared Resource at Huntsman Cancer Institute at the University of Utah and was supported by the National Cancer Institute of the National Institutes of Health under Award Number P30CA042014. The content is solely the responsibility of the authors and does not necessarily represent the official views of the NIH.

## Author contributions

Conception and design of the study: RDB, HL, CL, and SEG. Acquisition of data: HL, CL, ES, AL, and SEG. Analysis and interpretation of data: RDB, HL, CL, and SEG. Drafting or revising the manuscript: RDB, HL, CL, JW, BL, and SEG. All authors have approved the final article.

## Competing interests

HL, CL, ES, JW, AL, SEG, BL, and RDB have no competing interests.

## Materials and Methods

### Experimental Overview

ZNF865 was investigated to characterize its function, which at the time of writing is currently undefined, across a broad range of cell types and also investigated as a potential cell engineering tool for applications to cell therapy, tissue engineering, and gene therapy. CRISPR-guided gene modulation was used to investigate basic molecular processes and functions regulated by ZNF865 with RNAseq analysis as the starting point. Targeted gRNAs were designed for up- and down-regulation of ZNF865 and used to probe ZNF865’s role in regulating cell cycle, DNA replication, protein processing, RNA transport, and cellular senescence, which were indicated as the primary processes regulated in the RNAseq analysis, in human adipose-derived stem cells (ASCs), HEK 293 cells, Jurkat cells, and primary human nucleus pulposus cells. As a proof of concept, ZNF865 was then investigated for its ability to boost cell engineering outcomes related to tissue engineering and immune therapy and alter human senescent cell populations for potential senescence targeting gene therapies.

### General Cell Culture

All cell culture was performed in standard culture conditions (21% O_2_, 5% CO_2_, 37°C), with media changes every 2-3 days. *HEK 293*: Complete growth medium for cell culture for HEK 293 (ATCC CRL-1573) cells consists of HG-DMEM (ThermoFisher Scientific, Waltham, MA) supplemented with 10% fetal bovine serum (FBS) (ThermoFisher Scientific), 25μg/mL gentamicin (Corning), and 25mM HEPES (ThermoFisher). *hASC:* Complete growth medium for culturing of human adipose derived stem cells (ASCs, ATCC SCRC-4000) consists of Lonza ADSC Basal Medium (Lonza, Lexington, MA PT-3273), 10% MSC FBS (ThermoFisher Scientific), 5mL GlutaMax (ThermoFisher Scientific), 30μg/mL Gentamicin (Corning), and 15ng/mL Amphotericin (ThermoFisher Scientific). *Jurkat:* Complete growth medium for cell culture of Jurkat (ATCC TIB-152) cells consists of Advanced RPMI 1640 (ThermoFisher Scientific) supplemented with 10% FBS (ThermoFisher Scientific), 10mM HEPES (ThermoFisher Scientific), and 100 U/mL penicillin and 100μg/mL streptomycin (ThermoFisher Scientific). *hNPCs:* Primary human nucleus pulposus cells (hNPCs) were cultured in complete growth medium containing DMEM-HG (ThermoFisher Scientific) supplemented with 10% FBS (ThermoFisher Scientific), 50μg/mL gentamicin (Corning), 25mM HEPES (ThermoFisher Scientific), and 2ng/mL of recombinant human fibroblast growth factor-basic (rhFGF, Peprotech, Cranbury, NJ, 100-18C).

### Harvesting of Primary hNPCs and Culture

Human Healthy and Degenerative Nucleus pulposus (NP) tissue was obtained from surgical tissue waste and human nucleus pulposus cell (hNPC) isolation was performed as previously described^38^. Healthy hNPC tissue was obtained from trauma patients while degenerative hNPC tissue was obtained from patients undergoing discectomy. Briefly, NP tissue was rinsed twice with washing medium (DMEM-HG, 165 μg/mL gentamicin sulfate, 100μg/mL kanamycin sulfate, 1.25μg/mL amphotericin), minced, and enzymatically digested in washing medium with 0.3% (w/v) collagenase type II (Worthington Biochemical), 0.2% pronase (Sigma), and 5% FBS (ThermoFisher Scientific) for 2-3 hours at 37°C with 5% CO_2_ under gentle agitation^38^.

Isolated cells were passed through a 70μm cell strainer and washed twice. Cells were counted and plated at a density of 10,000 cells/cm^2^ in NP cell culture medium (DMEM-HG with 10% FBS, 50μg/mL gentamicin, 25mM HEPES) supplemented with fresh 2ng/mL fibroblast growth factor-2 (FGF-2) (Peprotech). Cells were cultured in this medium at 37°C and 5% CO_2_ in a humidified atmosphere and subcultured to 90% confluency, as previously described^38^.

### CRISPRi/CRISPRa

#### gRNA Design

Upregulation gRNA design was performed using Genome Target Scan 2 and ChopChop ^39,40^. The 5’-UTR, the promoter region up to 500bp upstream of the ZNF865 transcription start site (TSS), and up to 500bp upstream of the second exon were analyzed for gRNAs. The top 6 gRNAs from each analysis tool, with the least number of off-target sites, were compared and inspected using the BLAT tool of the UCSC genome browser to ensure minimal overlap between gRNAs and optimal placement near DNase hypersensitive regions^41^. A total of 6 gRNAS were selected for validation *in vitro*, as well as a gRNA that does not target the human genome (nontarget control/NTC). Oligos for ZNF865 gRNAs were synthesized by the University of Utah DNA/Peptide Synthesis core and ThermoFisher Scientific.

Downregulation gRNA design was performed as described above, using Genome Target Scan 2 (GT-Scan2) and ChopChop^39,40^. Briefly, a panel of 5 gRNAs were designed within the 5’-UTR and the promoter region up to 1000bp upstream of the ZNF865 TSS. Oligos for ZNF865 gRNAs were synthesized by the University of Utah DNA/Peptide Synthesis core and ThermoFisher Scientific.

#### Cloning

gRNAs, and a nontarget control (NTC) that does not target the human genome, were synthesized, annealed, phosphorylated, and ligated into individual U6-gRNA-UbC-DsRed-P2A-Bsr (upregulation gRNA) lentiviral upregulation expression vectors (Addgene, #83919) or KRAB CRISPRi downregulation expression vector pLV-hUbC-dCas9-KRAB-T2A-GFP (Addgene, #67620). Successful gRNA insertion was verified through Sanger sequencing^12,29,38^.

**Table 1:**
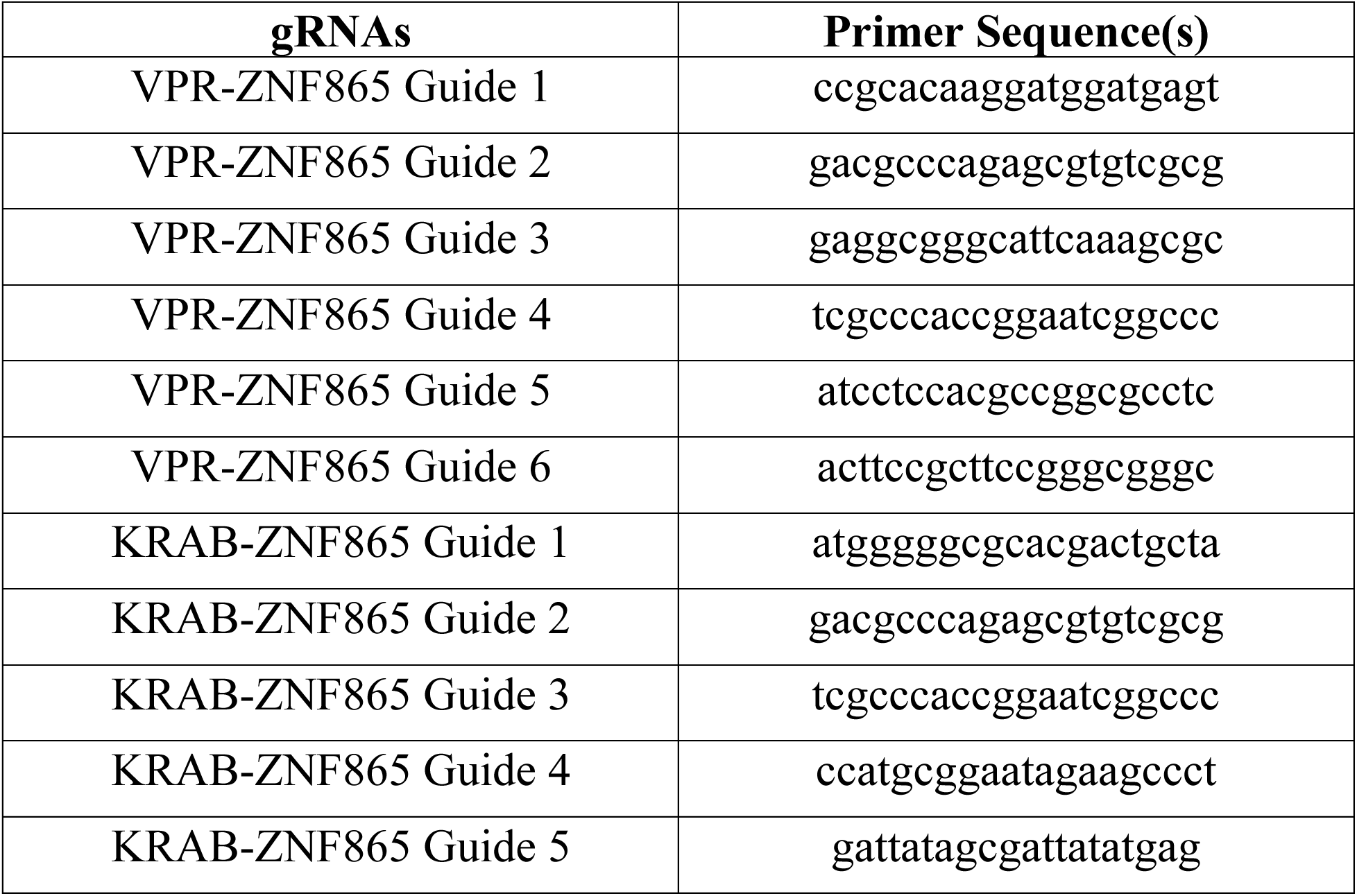
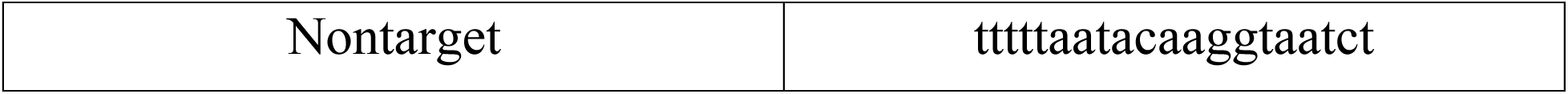
sgRNA primers for targeting the CRISPR system to ZNF865 and a nontarget control for upregulation and downregulation.

#### Lentivirus Production

The gRNA plasmid DNA was used to produce lentivirus, as previously described^12,29,38^. The amplified gRNA plasmid DNA was co-transfected into HEK 293T cells with psPAX2 (Addgene, plasmid #12260) and pMD2.G (Addgene, plasmid #12259) lentiviral packaging plasmids to create a lentivirus, as previously described^38^. Briefly, HEK 293T cells were seeded at a density of 62,500 cells/cm^2^. The following day, the lentiviral plasmids were added to the cells with Lipofectamine 2000 (ThermoFisher Scientific), following the manufacturer’s protocol. After 24 hours, the cell supernatant was discarded and replaced with fresh medium. Cell supernatant containing the lentiviral vectors was collected at 48 and 72 hours, concentrated to 100X, and stored at -80°C until use^29^.

#### CRISPRa Upregulation

##### dCas9-VPR Transduction

*HEK 293/ASC:* HEK 293 cells and ASCs were plated at a density of 20,000 cells/cm^2^ and 5,000 cells/cm^2^, respectively, allowed to attach overnight, and transduced with EF-1α-dCas9-VPR-PuroR (dCas9-VPR) lentivirus, (1:20 dilution) in growth medium containing 4μg/mL polybrene, the following day. Both cell types were subjected to antibiotic selection (1μg/mL) over the course of 3 days to select for dCas9-VPR expressing cells. *Jurkat:* Jurkat cells were suspended in media at a density of 50,000 cells/mL and transduced with EF-1α-dCas9-VPR-eGFP (dCas9-VPR) lentivirus, (1:20 dilution) in growth medium containing 4μg/mL, the following day. 72 hours after transduction, cells were subjected to fluorescence activated cell sorting (FACS) to select for GFP+ expressing cells. *hNPCs:* hNPCs (p4) were plated at a density of 20,000 cells/cm^2^, allowed to attach overnight, and transduced with EF-1a-dCas9-VPR-eGFP lentivirus, (1:2 dilution) in growth medium containing 4μg/mL, the following day. 72-hours following transduction hNPCs were subjected to FACS to select for GFP+ expressing cells.

##### gRNA Transduction

After dCas9-VPR transduction, dCas9-VPR expressing HEK 293 cells, hNPCs, and ASCs were transduced with gRNA targeting lentiviral vectors. *HEK293-VPR:* HEK 293-VPR cells were plated at a density of 20,000 cells/cm^2^ and transduced with gRNA vector (1:250 dilution) targeting ZNF865 or non-target the following day in complete media. *hNPC-VPR*: hNPCs were plated at a density of 5,000 cells/cm^2^ in a 24-well plate and transduced with gRNA vector (1:160 dilution) targeting ZNF865 or non-target in complete media the following day. *Jurkat-VPR*: Jurkat cells expressing dCas9-VPR were suspended in a 24-well plate at a density of 50,000 cells/mL and transduced the following day (1:160 dilution) with gRNA vector targeting ZNF865 or non-target in complete media. *ASC-VPR*: hNPCs were plated at a density of 5,000 cells/cm^2^ in a 24-well plate and transduced with gRNA vectors (1:160 dilution) targeting ZNF865, ZNF865/COL2A1/ACAN, or non-target in complete media the following day. Transduced HEK 293 and ASCs were examined for fluorescence after 48 hours and showed near 100% transduction efficiency. gRNA Transduced dCas9-VPR expressing Jurkat cells and hNPCs were subjected to FACS to select for successfully transduced cells and resuspended or plated at their appropriate density.

#### CRISPRi Downregulation

##### gRNA Transduction

For ZNF865 downregulation, naïve HEK-293, naïve ASCs, and naïve hNPCs were plated at a density of 20,000 cells/cm^2^, 5,000 cells/cm^2^, or 10,000 cells/cm^2^ respectively, in 24-well plates and allowed to attach overnight^38^. The following day, 100X sgRNA targeting ZNF865 or non-target virus was diluted 1:16 in complete growth medium supplemented with 4μg/mL polybrene and used to transduce cells. Cells were examined for fluorescence after 72-hours and showed near 100% transduction efficiency.

## ZNF865 Characterization

### ZNF865 qRT-PCR

Successfully transduced cells were analyzed for changes in ZNF865 gene expression by qRT-PCR (n=4). 72-hours post-transduction, RNA was isolated and harvested using the Quick-RNA Micro Kit (Zymo Research, R1051), complementary DNA (cDNA) was synthesized from the purified RNA with high-capacity cDNA reverse transcription kit with RNAse inhibitor (Applied Biosystems). cDNA was then used for qRT-PCR with TaqMan gene expression assays (ThermoFisher) for ZNF865 (Hs05052648_s1). Beta-2-microglobulin (B2M, Hs00187842_m1) was used as an internal standard and changes in ZNF865 expression was normalized to B2M expression^42^. Fold-change in mRNA expression relative to the NTC or ZNF865 edited cells was calculated using the ΔCT method.

### RNA-Sequencing

RNA-sequencing (RNAseq) was utilized to evaluate differential gene expression due to ZNF865 upregulation. HEK 293 cells, ASCs, and ACAN/COL2A1 ASCs were analyzed for global changes in gene expression after modification with ZNF865 or NTC CRISPRa vectors. Briefly, Total RNA was isolated from samples using a Quick-RNA Kit (Zymo Research, Irvine, CA). Isolated Total RNA (100-1000ng) was prepared using an Illumina TruSeq Stranded mRNA Library Prep Kit with PolyA Selection and samples were submitted to the High-Throughput Genomics Shared Resource Core at the Huntsman Cancer Institute for sequencing on a NovaSeq 6000, utilizing a NovaSeq Reagent Kit v1.5_150×150bp sequencing with 33 million reads per sample. Using previously described methods, data was normalized and compared to non-target control cells^12,38^.

Sequencing reads were aligned to hg38 build of the human genome and reads mapping UCSC known genes were counted using featureCounts from the SubRead package^43,44^. Reads were normalized and differential analysis was performed using DESeq2^43,45^. Enriched GO biological and molecular processes were determined from significantly regulated genes using Enrichr^46–48^. Comprehensive gene expression profiling was performed using NetworkAnalyst to evaluate the transcription factor-gene interactions with the common upregulated genes across all three cell types^19,20^.

### Cell Proliferation Quantification

Following successful transduction, HEK 293, ASCs, hNPCs, and Jurkat cells transduced with either the CRISPRa upregulation system or CRISPRi downregulation system (n=4) were evaluated for cell proliferation over the course of 3-10 days. Individual cells were counted using ImageJ or a hemacytometer^49^. Briefly, fluorescing cell counts were obtained by uploading images into a stack and thresholding the images to ensure only individual cells are shown within the image. After thresholding, cell counts were obtained by analyzing particles, generating a mask, and then counting the masks generated while excluding cells on the edges of the image. In instances where cell density was too great for thresholding and visually isolating individual cells, cells were manually counted in ImageJ using the cell counter plugin^49^.

### Cell Cycle Analysis

For upregulation cell cycle analysis, ZNF865 and NTC edited cells were grown to confluency in a T-75 flask, lifted from the flask, and resuspended in growth medium at a concentration of 1 million cells/mL. Following the manufacturer’s instructions, 2 drops of Vybrant™ DyeCycle™ Violet Ready Flow™ Reagent (ThermoFisher Scientific, R37172) was added to the suspension before being incubated at 37°C for 30 minutes. Following incubation, cells were analyzed on a Cytoflex S Flow Cytometer (Beckman Coulter Life Sciences, Indianapolis, IN). Flow cytometry data was analyzed using FlowJo Flow Cytometry Software (BD Biosciences). Gates for selecting individual cells, dsRed expressing cells, and DyeCycle™ violet expressing cells were used to isolate and analyze our cells of interest.

For downregulation cell cycle analysis, HEK 293 cells were plated in a 6-well plate at a density of 50,000 cells/mL. The following day, gRNA virus was diluted 1:25 in complete growth medium supplemented with 4μg/mL polybrene and used to transduce cells. All transduced cells were examined for fluorescence after 48 hours and showed near 100% transduction efficiency. 72-hours after transduction, cells were lifted and resuspended in growth medium at a concentration of 1 million cells/mL and cell cycle was analyzed as described above.

### Cellular Senescence Analysis (SA-β-galactosidase, p16^INK4A^, p21^CIP1^)

#### SA-β-Galactosidase

Primary hNPCs were plated in a 24-well plate at a concentration of 10,000 cells/cm^2^ and allowed to attach overnight. The following day attached cells were transduced with the ZNF865 (Guide 2 and Guide 3) or NTC downregulation expression cassettes. Cells were cultured for 10 or 24 days, at which point they were fixed and stained for SA-β-galactosidase using a Senescent Cell Histochemical Staining Kit (Sigma-Aldrich, CS0030-1KT). Cellular senescence was quantified by counting SA-β-galactosidase stained and unstained cells using ImageJ.

#### P16^INK4A^ and p21^CIP1^Immunofluorescence

Primary hNPCs were plated into cell culture chamber slides at a concentration of 10,000 cells/cm^2^ and then transduced with the ZNF865 or NTC downregulation expression cassettes. Cells were then cultured for 10 or 24 days after which the cells were fixed with 10% formalin. Following fixation cells were permeabilized with 0.2% Triton X in PBS for 5 min. Cells were then washed three times with PBS and treated with blocking solution (5% goat serum [MP Biomedicals] in PBS for 1 hour at room temperature). Following treatment with blocking solution, the blocking solution was removed, and the cells were treated with p16^INK4A^ (p16) antibody (1:100 in 1% BSA in PBS; Proteintech) or p21^CIP1^ (p21) antibody (1:100 in 1% BSA in PBS; Proteintech), normal rabbit IgG (Invitrogen) or BSA only negative controls and incubated at room temperature for 2 hours.

Following incubation cells were washed three times with PBS, treated with goat anti-rabbit Coralite 594 (1:100 Proteintech) in 1% BSA in PBS, and incubated for 1 hour at room temperature. Subsequently, cells were rinsed three times with PBS, treated with DAPI solution (3ng/mL) for 10 minutes, and mounted with Vectamount Mounting Medium prior to fluorescence imaging (Leica SP8 Confocal).

## Cell Engineering

### Rescuing Cell Populations from Senescence

Degenerative primary dCas9-VPR expressing hNPCs were plated in a 24-well plate at a concentration of 10,000 cells/cm^2^ and allowed to attach overnight. The following day cells were transduced with the ZNF865 (Guide 1) upregulation lentivirus, as described previously in *gRNA transduction.* Successfully transduced cells were cultured for 10 or 24 days, at which point they were fixed and stained for SA-β-galactosidase, p16^INK4A^, and p21^CIP1^, as described previously in *Cellular Senescence Analysis*.

### Jurkat T-Cell Activation

VPR-ZNF865 and -NTC expressing Jurkat cells were plated in a 24-well plate at a density of 250,000 cells/mL (n=5). The following day, T-cells were stimulated using with 1X Concanavalin A (00-4978-93, ThermoFisher Scientific). Following 3 days of incubation, T-cell proliferation was evaluated using CCK8, following the manufacturers protocol, and T-cell supernatant was harvested and analyzed for IL-2 (900-M12, ThermoFisher Scientific) and IFN-γ (900-M27, ThermoFisher Scientific) cytokine secretion using an Enzyme-linked immunosorbent assay (ELISA) following the manufacturer’s protocol.

### Pellet Culture of hASCs

To evaluate extracellular matrix (ECM) deposition in our ACAN/COL2A1 edited cells, 3D pellet culture analysis was performed as previously described^12,29^. Briefly, pellets were formed by resuspending ACAN/COL2A1-NTC and ACAN/COL2A1-ZNF865 edited hASCs at a concentration of 1.25 million cells/mL in a serum-free growth medium and 200μL aliquots of the cell suspension was pipetted into individual wells of a 96-well u-bottom plate and spun at 270G for 5 minutes at 4°C. Cells were allowed to contract overnight and form pellets. The following day pellets were gently lifted from the bottom of the plate. Media was changed on the pellets every 2-3 days for 21-days and supernatant was collected during every media change. After 21-days pellets were harvested and either papain digested for biochemical analysis or fixed in formalin and submitted for histological analysis, as previously described^12,29^.

### Pellet Biochemical Analysis

The total amount of collagen content contained within papain digested pellets and supernatant (n=10) was analyzed using a modified hydroxyproline assay^12,29^. Hydroxyproline assay was performed as previously described, with the same adjustments for hydrolysis time and adjusting the pH of the oxidation buffer to 6.5 with glacial acetic acid^12,29^. The total amount of DNA content and sGAG content within papain digested pellets and supernatant (n=10) was analyzed using a previously described Hoescht dye assay and a dimethylmethylene blue (DMMB) assay^12,29,50,51^.

### Macroscopic Pellet Imaging and Size Analysis

After 21-days of pellet culture, pellets were imaged (Canon Rebel T3) in their respective wells for qualitative comparison of gross pellet morphology, as previously described^12,29^. Volume was estimated by assuming the pellets were spheres, measuring the average diameter across the pellet, and calculating volume (n=15).

### Pellet Culture Histological Analysis

To prepare pellets for staining, pellets (n=5) were fixed in a 10% neutral-buffered formalin solution for 24-hours, embedded in paraffin, and 5μm sections were mounted on glass slides^12,29^. Sections for each sample were stained with Alcian blue (pH 2.5; Newcomer Supply) and counterstained in Nuclear-fast red (Newcomer Supply). Briefly, slides were deparaffinized and rehydrated to distilled water, suspended in 3% acetic acid for 3 minutes, suspended in Alcian blue solution at room temperature for 30 minutes, washed in distilled water for 2 minutes, suspended in Nuclear-fast red solution for 5 minutes, washed in tap water, dehydrated, cleared, and cover slipped.

### DAPS Cell Seeding and Culture

Naïve ASCs, ACAN/COL2A1-NTC, and ACAN/COL2A1-ZNF865 edited ASCs were evaluated for cartilage deposition *in vitro* in DAPS over 5-weeks of culture. DAPS were fabricated to be 10mm in diameter and 3mm in height, seeded, and cultured as previously described^16^. Briefly, the NP region of DAPS was formed by suspending ASCs in chemically defined media at a density of 40 x 10^6^ cells/mL mixing with molten 4% w/v agarose (49°C, Type VII, Sigma-Aldrich), and cast into 6-well plates to generate agarose slabs at a final density of 20 x 20^6^ cells/mL and 2% w/v agarose gel. Sterilized biopsy punches generated NP regions of DAPS that were 3mm thick and 5mm in diameter. NP regions were cultured in isolation for 2.5 weeks on an orbital shaker prior to combining with AF regions of DAPS. Full DAPS culture was performed in chemically defined media consisting of: high glucose DMEM supplemented with 1% PSF, 40 ng/mL dexamethasone (Sigma-Aldrich), 50 μg/mL ascorbate 2-phosphate (Sigma-Aldrich), 40 μg/mL L-proline (Sigma-Aldrich), 100 μg/mL sodium pyruvate (Corning Life Sciences, Corning, NY), 0.1% insulin, transferrin, and selenious acid (ITS Premix Universal Culture Supplement; Corning), 1.25 mg/mL bovine serum albumin (Sigma-Aldrich), 5.35 μg/mL linoleic acid (Sigma-Aldrich), and 10 ng/mL TGF-β3 (R&D Systems, Minneapolis, MN) with media changes every 2-3 days, with each hydrogel receiving 1.5mL of media per change.

AF regions of DAPS were fabricated using a 14.3% w/v solution of poly(ε-caprolactone) (PCL) dissolved in a 1:1 mixture of tetrahydrofuran and N,N-dimethylformamide and electrospun onto a grounded rotating mandrel, as previously described^52^. Native lamellar architecture was replicated by fabricating PCL sheets that were 250μm thick and were cut at a 30° angle to orient native fiber direction. Sheets were cut into 3mm thick and 150mm in length strips which were hydrated through a gradient of ethanol, before coating strips overnight with 20μg/mL of fibronectin (Sigma-Aldrich) in PBS. ASCs were suspended in growth medium and seeded onto PCL strips at a density of 1.5 x 10^6^ cells per side and cultured for 1-week. Cell-seeded strips of AF region were assembled as strips 2 strips thick with opposing fiber directions (± 30°) prior to wrapping the strips around a custom mold and cultured on an orbital shaker, as previously described^37^. Following 1.5-weeks of culture around the mold, AF regions were removed from the mold and combined with NP regions, at which point combined NP and AF regions of DAPS were cultured for 2.5-weeks, resulting in a total of 5-weeks of culture. AF regions were cultured in the same chemically defined media as NP regions with media changes every 2-3 days.

DAPS were evaluated statistically by a Pearson’s Chi-Squared Analysis with DAPS either being considered Success or Fail. Successful DAPS were DAPS that produced sufficient ECM to maintain shape and not have the AF region begin to unroll. Failed DAPS were DAPS that did not maintain shape, AF began to unroll, and had to be pinned together during the 5-week culture period.

### DAPS Histological Evaluation

For histological assessment of matrix deposition after 5-weeks of culture, DAPS were fixed in 10% neutral buffered formalin, embedded in paraffin, and sectioned in the axial plane to 10μm thickness. Sections were stained for sGAG deposition using Alcian blue and/or collagen deposition using picrosirius red.

### Statistics

RNA-seq statistical analysis was performed and described in their respective section (Enrichr, DESEQ2, α=0.05). Statistical analysis of qRT-PCR, proliferation, and biochemical data was performed using JMP Pro using a one-way analysis of variance (ANOVA) with a Tukey’s post-hoc analysis (α=0.05 for all tests).

## Notes

### Competing Interest Statement

The authors have declared no competing interest.

## References Cited

1. Laity, J. H., Lee, B. M. & Wright, P. E. Zinc finger proteins: new insights into structural and functional diversity. Curr Opin Struct Biol 11, 39–46 (2001).

2. Wolfe, S. A., Nekludova, L. & Pabo, C. O. DNA RECOGNITION BY Cys 2 His 2 ZINC FINGER PROTEINS. www.annualreviews.org (2000).

3. Singh, J. K. et al. Zinc finger protein ZNF384 is an adaptor of Ku to DNA during classical non-homologous end-joining. Nat Commun 12, (2021).

4. Ghaleb, A. M. & Yang, V. W. Krüppel-like factor 4 (KLF4): What we currently know. Gene vol. 611 27–37 Preprint at 10.1016/j.gene.2017.02.025 (2017).

5. Stubbs, L., Sun, Y. & Caetano-Anolles, D. Function and Evolution of C2H2 Zinc Finger Arrays. Subcell Biochem 52, 75–94 (2011).

6. Bateman, A. et al. UniProt: the Universal Protein Knowledgebase in 2023. Nucleic Acids Res 51, D523– D531 (2023).

7. Frietze, S. & Farnham, P. J. Transcription factor effector domains. Subcell Biochem 52, 261–77 (2011).

8. Hong, K. et al. Comprehensive analysis of ZNF family genes in prognosis, immunity, and treatment of esophageal cancer. BMC Cancer 23, 301 (2023).

9. Gilbert, L. A. et al. Genome-Scale CRISPR-Mediated Control of Gene Repression and Activation. Cell 159, 647–61 (2014).

10. Horlbeck, M. A. et al. Compact and highly active next-generation libraries for CRISPR-mediated gene repression and activation. (2016) doi:10.7554/eLife.19760.001.

11. Levis, H. et al. Multiplex gene editing to promote cell survival using low-pH clustered regularly interspaced short palindromic repeats activation (CRISPRa) gene perturbation. Cytotherapy (2023) doi:10.1016/j.jcyt.2023.05.001.

12. Farhang, N. et al. Synergistic CRISPRa-regulated chondrogenic extracellular matrix deposition without exogenous growth factors. Tissue Eng Part A 26, 1169–1179 (2020).

13. Chavez, A., et al. Highly efficient Cas9-mediated transcriptional programming. communicaHons 326, (2015).

14. Thakore, P. I. et al. highly specific epigenome editing by crisPr-cas9 repressors for silencing of distal regulatory elements. (2015) doi:10.1038/nmeth.3630.

15. Gullbrand, S. E., et al. Long-term mechanical funcHon and integraHon of an implanted Hssue-engineered intervertebral disc. Sci. Transl. Med vol. 10 http://stm.sciencemag.org/ (2018).

16. Gullbrand, S. E. et al. Towards the scale up of tissue engineered intervertebral discs for clinical application. Acta Biomater 70, 154–164 (2018).

17. Kanehisa, M. KEGG: Kyoto Encyclopedia of Genes and Genomes. Nucleic Acids Res 28, 27–30 (2000).

18. Najafabadi, H. S. et al. C2H2 zinc finger proteins greatly expand the human regulatory lexicon. Nat Biotechnol 33, 555–62 (2015).

19. Xia, J., Gill, E. E. & Hancock, R. E. W. NetworkAnalyst for statistical, visual and network-based meta-analysis of gene expression data. Nat Protoc 10, 823–44 (2015).

20. Zhou, G. et al. NetworkAnalyst 3.0: a visual analytics plamorm for comprehensive gene expression profiling and meta-analysis. Nucleic Acids Res 47, W234–W241 (2019).

21. Kent, W. J. et al. The Human Genome Browser at UCSC. Genome Res 12, 996–1006 (2002).

22. Heathman, T. R. et al. The translation of cell-based therapies: Clinical landscape and manufacturing challenges. RegeneraHve Medicine vol. 10 49–64 Preprint at 10.2217/rme.14.73 (2015).

23. Thakore, P. I. et al. Highly specific epigenome editing by CRISPR-Cas9 repressors for silencing of distal regulatory elements. Nat Methods 12, 1143–9 (2015).

24. Silwal, P. et al. Cellular Senescence in Intervertebral Disc Aging and Degeneration: Molecular Mechanisms and Potential Therapeutic Opportunities. Biomolecules 13, 686 (2023).

25. Patil, P. et al. Cellular senescence in intervertebral disc aging and degeneration. Curr Mol Biol Rep 4, 180–190 (2018).

26. Noren Hooten, N. & Evans, M. K. Techniques to Induce and Quantify Cellular Senescence. J Vis Exp (2017) doi:10.3791/55533.

27. Rayess, H., Wang, M. B. & Srivatsan, E. S. Cellular senescence and tumor suppressor gene p16. Int J Cancer 130, 1715–1725 (2012).

28. Watanabe, S., Kawamoto, S., Ohtani, N. & Hara, E. Impact of senescence-associated secretory phenotype and its potential as a therapeutic target for senescence-associated diseases. Cancer Science vol. 108 563–569 Preprint at haps://doi.org/10.1111/cas.13184 (2017).

29. Farhang, N. et al. CRISPR-Based Epigenome Editing of Cytokine Receptors for the Promotion of Cell Survival and Tissue Deposition in Inflammatory Environments. doi:10.1089/ten.tea.2016.0441.

30. Collado, M., Blasco, M. A. & Serrano, M. Cellular Senescence in Cancer and Aging. Cell 130, 223–233 (2007).

31. Price, J. S. et al. The role of chondrocyte senescence in osteoarthritis. Aging Cell 1, 57–65 (2002).

32. Bhat, R. et al. Astrocyte Senescence as a Component of Alzheimer’s Disease. PLoS One 7, e45069 (2012).

33. Sadelain, M., Rivière, I. & Riddell, S. Therapeutic T cell engineering. Nature vol. 545 423–431 Preprint at10.1038/nature22395 (2017).

34. Bowles, R. D. & Seaon, L. A. Biomaterials for intervertebral disc regeneration and repair. Biomaterials 129, 54–67 (2017).

35. GBD 2017 Disease and Injury Incidence and Prevalence Collaborators. Global, regional, and national incidence, prevalence, and years lived with disability for 354 diseases and injuries for 195 countries and territories, 1990-2017: a systematic analysis for the Global Burden of Disease Study 2017. Lancet 392, 1789–1858 (2018).

36. O’Halloran, D. M. & Pandit, A. S. Tissue-engineering approach to regenerating the intervertebral disc. Tissue Eng 13, 1927–54 (2007).

37. Nerurkar, N. L., Sen, S., Huang, A. H., Ellioa, D. M. & Mauck, R. L. Engineered disc-like angle-ply structures for intervertebral disc replacement. Spine (Phila Pa 1976) 35, 867–73 (2010).

38. Farhang, N. et al. Lentiviral CRISPR Epigenome Editing of Inflammatory Receptors as a Gene Therapy Strategy for Disc Degeneration. Hum Gene Ther 30, 1161–1175 (2019).

39. O’Brien, A. & Bailey, T. L. GT-Scan: identifying unique genomic targets. BioinformaHcs 30, 2673–2675 (2014).

40. Labun, K. et al. CHOPCHOP v3: expanding the CRISPR web toolbox beyond genome editing. Nucleic Acids Res 47, W171–W174 (2019).

41. Kent, W. J. BLAT -- The BLAST-like alignment tool. Genome Res 12, 656–664 (2002).

42. Vandesompele, J. et al. Accurate normalizaHon of real-Hme quanHtaHve RT-PCR data by geometric averaging of mulHple internal control genes. hap://genomebiology.com/2002/3/7/research/0034.1Correspondence:.rankSpeleman. (2002).

43. Kim, D., Langmead, B. & Salzberg, S. L. hisAt: a fast spliced aligner with low memory requirements. ArHcles nAture methods 12, 357 (2015).

44. Liao, Y., Smyth, G. K. & Shi, W. featureCounts: an efficient general purpose program for assigning sequence reads to genomic features. BioinformaHcs 30, 923–30 (2014).

45. Love, M. I., Huber, W. & Anders, S. Moderated estimation of fold change and dispersion for RNA-seq data with DESeq2. Genome Biol 15, 550 (2014).

46. Kuleshov, M. V et al. Enrichr: a comprehensive gene set enrichment analysis web server 2016 update. Nucleic Acids Res 44, (2016).

47. Chen, E. Y. et al. Enrichr: interacHve and collaboraHve HTML5 gene list enrichment analysis tool. http://amp.pharm.mssm.edu/Enrichr. (2013).

48. The Gene Ontology project in 2008. doi:10.1093/nar/gkm883.

49. Schneider, C. A., Rasband, W. S. & Eliceiri, K. W. NIH Image to ImageJ: 25 years of image analysis. Nat Methods 9, 671–5 (2012).

50. Zheng, C. & Levenston, M. Fact versus artifact: Avoiding erroneous estimates of sulfated glycosaminoglycan content using the dimethylmethylene blue colorimetric assay for tissue-engineered constructs. Eur Cell Mater 29, 224–236 (2015).

51. Kim, Y.-J., Sah, R. L. Y., Doong, J.-Y. H. & Grodzinsky, A. J. Fluorometric assay of DNA in cartilage explants using Hoechst 33258. Anal Biochem 174, 168–176 (1988).

52. Nerurkar, N. L., Ellioa, D. M. & Mauck, R. L. Mechanics of oriented electrospun nanofibrous scaffolds for annulus fibrosus tissue engineering. J Orthop Res 25, 1018–28 (2007).

